# Automated landmark and semilandmark annotation for wing geometric morphometrics in Diptera using deep learning

**DOI:** 10.64898/2026.04.17.719146

**Authors:** Kristopher Nolte, Jan Baumbach, Philip Kollmannsberger, Felix Gregor Sauer, Renke Lühken

**Author notes:** shared last authors (Felix Gregor Sauer, Renke Lühken).

## Abstract

**1.** Diptera represent a diverse insect order, including vectors of human and animal pathogens. Their accurate species identification remains a major bottleneck in ecological and epidemiological studies. Morphological identification requires taxonomic expertise, while molecular methods are costly and not universally reliable. Wing geometric morphometrics offers an alternative, but manual landmark annotation is time-consuming and introduces observer bias. **2.** We developed ITHILDIN, an automated pipeline for landmark and semilandmark annotation of Diptera wings, combining UNet++ segmentation and an Hourglass landmark prediction model. Using mosquitoes as the primary model system, we extended an existing repository with 5,793 additional images. Models were trained on 5991 annotations of landmarks and segmentations and then evaluated on 12,522 images across 34 taxa. We assessed landmark prediction accuracy against human observers and ML-morph, evaluated species identification using Linear Discriminant Analysis on 17 homologous landmarks and 52 semilandmarks, and tested out-of-distribution generalisation by reproducing an independent study. Transferability was demonstrated by adapting the pipeline to the Dipteran families Drosophilidae and Glossinidae. **3.** The Hourglass model achieved a mean landmark error of 4.5 pixels (95% CI: 4.3–4.6), within human observer variability (4.7 pixels, 95% CI: 4.4–5.0) and substantially outperforming ML-Morph (12.7 pixels, 95% CI: 11.1–14.2). The semilandmark-based approach for species identification achieved 91% balanced accuracy across 34 taxa, comparable to CNN performance (94%). On out-of-distribution data, the landmark pipeline generalised substantially better than the CNN and a soft-voting ensemble of the landmark and CNN classifiers achieved 88% balanced accuracy on a replicated study. **4.** Combining geometric morphometrics with deep learning provides a reproducible, interpretable, and generalisable alternative to black-box CNN classifiers for Diptera wing analysis. By acting as a consistent single observer comparable to human annotation, the system eliminates inter-observer bias, enabling large-scale and cross-study morphometric analyses of Dipteran wings. The system is publicly available at www.ithildin.bnitm.de and transferable to other Diptera families with moderate retraining effort.

**Data availability:** Images used in this study are accessible under CC BY 4.0 license at https://doi.org/10.6019/S-BIAD1478. Downloadable and installable docker application can be accessed on the applications’ git page: https://anonymous.4open.science/r/ITHILDIN-4313/

## 1 Introduction

Diptera are one of the most diverse insect orders, with over 150,000 described species. They serve as pollinators, decomposers, and food sources, playing essential roles in ecosystem functioning (Courtney et al. 2017). The order also includes several families whose species are blood-sucking and are among the most important vectors of disease (Lehane and Lehane 2005). Climate change, urbanisation, and global trade are expanding the geographic range of these vectors, increasing transmission risk in previously unaffected regions (de Souza and Weaver 2024). Both for ecological research and for vector surveillance, accurate species identification is essential yet it remains a persistent bottleneck (ECDC 2021).

Morphological identification of Diptera requires trained taxonomic experts, a scarce resource globally (Wilkerson, Linton, and Strickman 2021). Molecular methods such as DNA barcoding offer an expert-independent alternative, but they are costly, time-consuming, and not universally reliable (Farlow, Russell, and Burkot 2020). Cryptic species complexes, incomplete reference databases, and degraded specimens frequently limit their utility as well (Piper et al. 2019).

Recent advances in computer vision offer potential solutions. Convolutional neural networks (CNNs) and transformer-based models can classify insect images with high accuracy (Couret et al. 2020; Goodwin et al. 2021; Sauer et al. 2024; Zhao et al. 2022). However, these approaches have significant limitations. They function as “black boxes” with limited interpretability (Nolte et al. 2024; Nolte, Baumbach, et al. 2025), are highly vulnerable to spurious correlations (Geirhos et al. 2020), generalise poorly to out-of-distribution data and are constrained to fixed classification tasks (D’Amour et al. 2022).

Wing geometric morphometrics provides a promising alternative. It captures the shape and size of the wings using the coordinates of homologous anatomical landmarks on the vein junctions. These landmark coordinates are aligned through Procrustes superimposition to remove the effects of size, position, and orientation and can then be used to quantify shape variation among specimens (Zelditch 2025). This has been demonstrated efficient for species classification in mosquitoes (Champakaew et al. 2021; Sauer et al. 2020) and other Diptera (Tatsuta, Takahashi, and Sakamaki 2018). Additionally, morphometric wing characters were successfully utilised to discriminate between cryptic species for which the morphological separation is difficult or impossible (Börstler et al. 2014). Beyond species identification, wing geometric morphometrics supports studies of intraspecies variation, asymmetry, wing size, allometry, and phenotypic plasticity (Lorenz et al. 2017). However, manual landmark annotation is time-consuming, introduces observer bias, and produces results that are often incompatible across studies (Dujardin, Kaba, and Henry 2010). These limitations have confined wing geometric morphometrics to small-scale, labour-intensive investigations.

Automated landmark annotation addresses these shortcomings. Rather than learning to classify images directly, a landmark-based system learns to locate anatomical points. This task is decoupled from downstream performance, i.e. classification accuracy, and is therefore less vulnerable to domain shift than CNN classifiers. Because the model is trained to predict anatomical landmarks rather than class labels, it must learn geometric structures that are necessary for accurate landmark localization, reducing the incentive to exploit spurious correlations such as lighting conditions or background features (Azulay and Weiss 2019; D’Amour et al. 2022). Additionally, this approach produces output that can be visually inspected and statistically validated.

The goal of automating insect wing landmark annotation has been pursued for over two decades. Houle et al. (2003) developed an automated wing measurement system for *Drosophila* based on manually designed feature extraction methods. However, this approach required highly standardised image acquisition and did not generalise to other species or imaging conditions. Subsequent refinements improved general performance but nevertheless they remained vulnerable to variation in image collection techniques (Eshghi et al. 2024; Nguyen et al. 2022; Rebelo et al. 2021). The advent of machine learning has accelerated progress considerably. More generalisable and retrainable tools such as ML-Morph developed by Porto and Voje (2020) were introduced. Convolutional models made feature extraction more flexible and faster to implement (Le et al. 2020). Geldenhuys et al. (2023) demonstrated that a segmentation-based approaches could automatically annotate landmarks on tsetse fly (Glossinidae) wings with high accuracy. However, in contrast to Drosophilidae where most current methods were developed for, most the wing veins of other dipteran families such as mosquitoes (Culicidae) or moth flies (Psychodidae) are covered with scales or hairs, which makes automated landmark annotation more challenging.

While deep learning–based landmark annotation for insect wings remains underdeveloped, progress has been made in related fields. Facial landmark detection and human pose estimation share the same objective of locating anatomical keypoints in images (Nguyen et al. 2022). Advances in these fields such as architectural innovations and improved loss functions, are directly transferable to entomological applications. Stacked Hourglass networks, originally developed for human pose estimation, capture multi-scale features and preserve local and global detail through repeated encoding and decoding (Newell, Yang, and Deng 2016). Boundary-aware guidance, as proposed by (Wu et al. 2018), directs model attention toward regions of interest by incorporating structural priors. An adaptive wing loss function further improves performance by differentially weighting foreground and background pixels, enhancing accuracy on fine anatomical structures (Wang, Bo, and Fuxin 2020). These methods provide the foundation for a robust landmark annotation system applicable to insect wings as demonstrated by Chumnoi et al. (2026).

Semilandmarks could even extend the approach of wing geometric morphometric analysis. Unlike “traditional” landmarks, which are limited to discrete points, semilandmarks capture shape variation along curves and surfaces (Gunz and Mitteroecker 2013). In wing geometric morphometrics, they can be placed along the veins, increasing the resolution of shape description (Chaiphongpachara and Laojun 2019). This is particularly valuable when distinguishing morphologically similar species or analysing subtle intraspecies variation (Farré et al. 2016). Automating this annotation process would largely eliminate the bottleneck of increased workload, making it feasible to place dense configurations of semilandmarks across large datasets.

Here, we present ITHILDIN (Integrated Tool for Homologous Image Landmark Detection in Insects using Neural networks), a fully automated system for landmark and semilandmark annotation of Diptera wings. We developed the system without relying on manually designed feature extraction or standardised imaging conditions. The pipeline combines deep learning–based segmentation and landmark prediction with downstream semilandmark placement, enabling robust annotation across diverse image sources. We used mosquitoes as the initial model system due to their morphological complexity, including complex venation and the presence of wing scales overlapping the veins and the availability of a large, annotated image repository, which we further extended (Nolte, Agboli, et al. 2025).

We evaluated ITHILDIN through four experiments. First, we compared landmark annotation performance to human annotators and current automated landmark systems, namely ML-morph (Porto and Voje 2020). Second, we assessed species identification performance across 34 mosquito taxa, comparing automated landmark-based classification using Linear Discriminant Analysis (LDA) with CNN-based approaches. Third, we tested out-of-distribution generalisation by reproducing an independent mosquito wing study (Jeon et al. 2024) using ITHILDIN. Lastly, we demonstrated cross-family transferability by adapting the system to the Dipteran families Drosophilidae and Glossinidae. A demonstrator of the system is publicly available at ithildin.bnitm.de and the complete system can be downloaded at https://anonymous.4open.science/r/ITHILDIN-4313/.

## 2 Materials and Methods

### 2.1 Overview Landmark Annotation Pipeline

IHTILDIN is a pipeline of processing steps and deep learning model predictions (Figure 1). It transforms a wing image into (semi)landmark coordinates and predicts the species. First, the image undergoes a subset of the preprocessing steps proposed by Nolte, Baumbach, et al. (2025). Here the background is removed, the wing horizontally aligned, and image features that could serve as spurious correlations are reduced using Contrast-Limited Adaptive Histogram Equalisation. Next, the processed wing image is segmented using an UNet++ model (Zhou et al. 2018). The segmentation and processed image are concatenated and passed into an Hourglass model, which predicts the landmark positions (Wu et al. 2018). Using the predicted landmarks and segmentation, a digital representation of the wing is constructed. This representation consists of landmarks as points and vein segments as paths. Semilandmarks are allocated at equidistant intervals along these veins. Additionally, a CNN trained according to Nolte, Baumbach et al. (2025), predicts the species based on the processed image. A decision module combines the CNN-based prediction with a landmark-based prediction from LDA to produce the final taxonomic classification.

**Figure 1:**
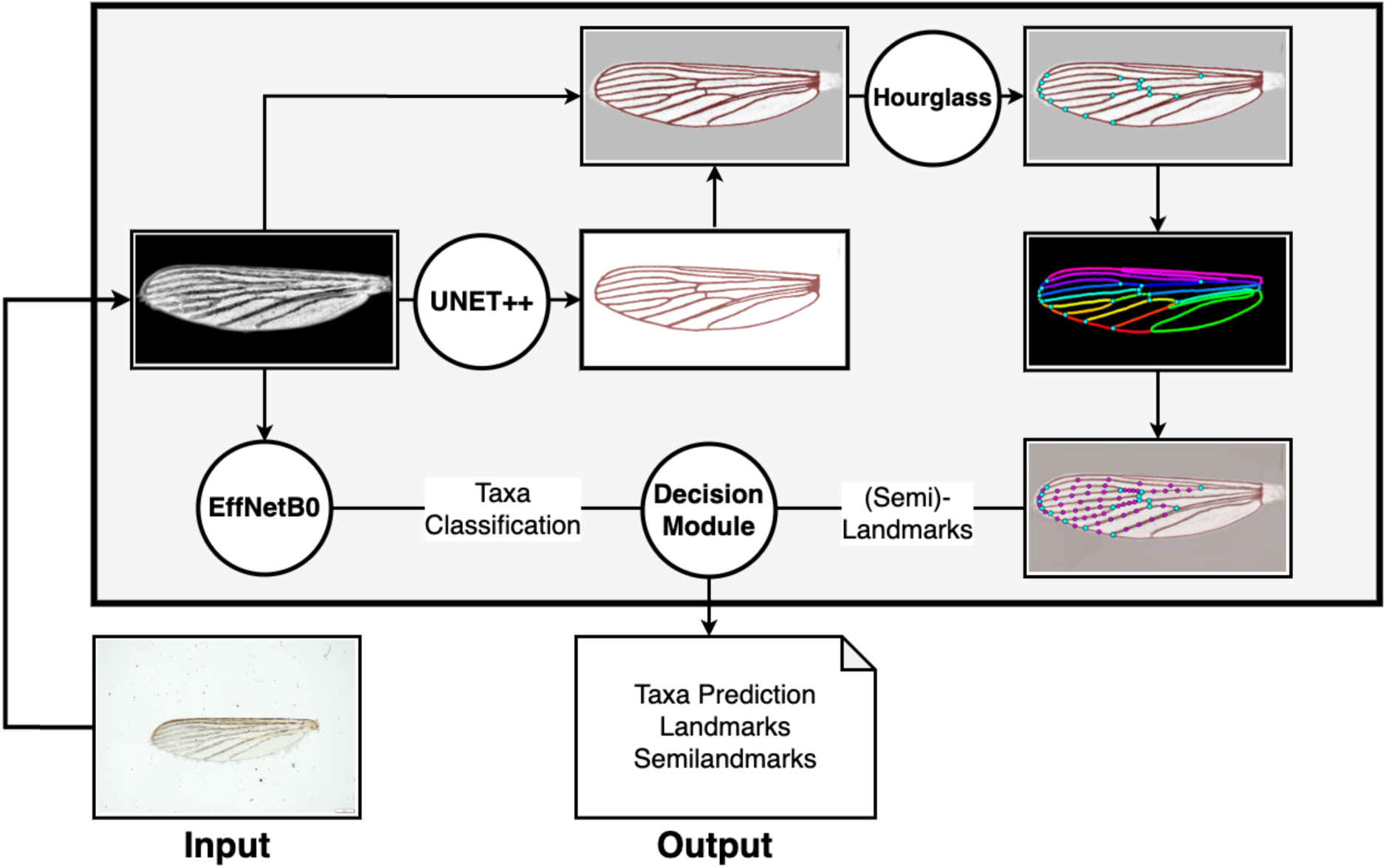
Schematic Workflow of ITHILDIN. Images are first pre-processed by the pipeline according to Nolte, Baumbach et al. (2025). Hereafter, a segmentation model (UNET++) extracts the vein pattern. The segmentation and the image are concatenated and fed into an Hourglass model which predicts the landmarks. Based on the landmark and the segmentation a representation of the wing is produced. This is used to place semilandmarks on the paths between the landmarks. Additionally, the pre-processed image is input in a CNN (ETNetB0) which predicts the species.

### 2.2 Pipeline Development

#### 2.2.1 Image Acquisition and Preprocessing

To train three computer vision models (UNet++, Hourglass, CNN), we used the mosquito wing image repository from Nolte, Agboli, et al. (2025). We sampled 15,621 undamaged wing images of this repository and expanded the dataset with additional collections (N = 5793), yielding 21,414 images (Table 1).

**Table 1:**
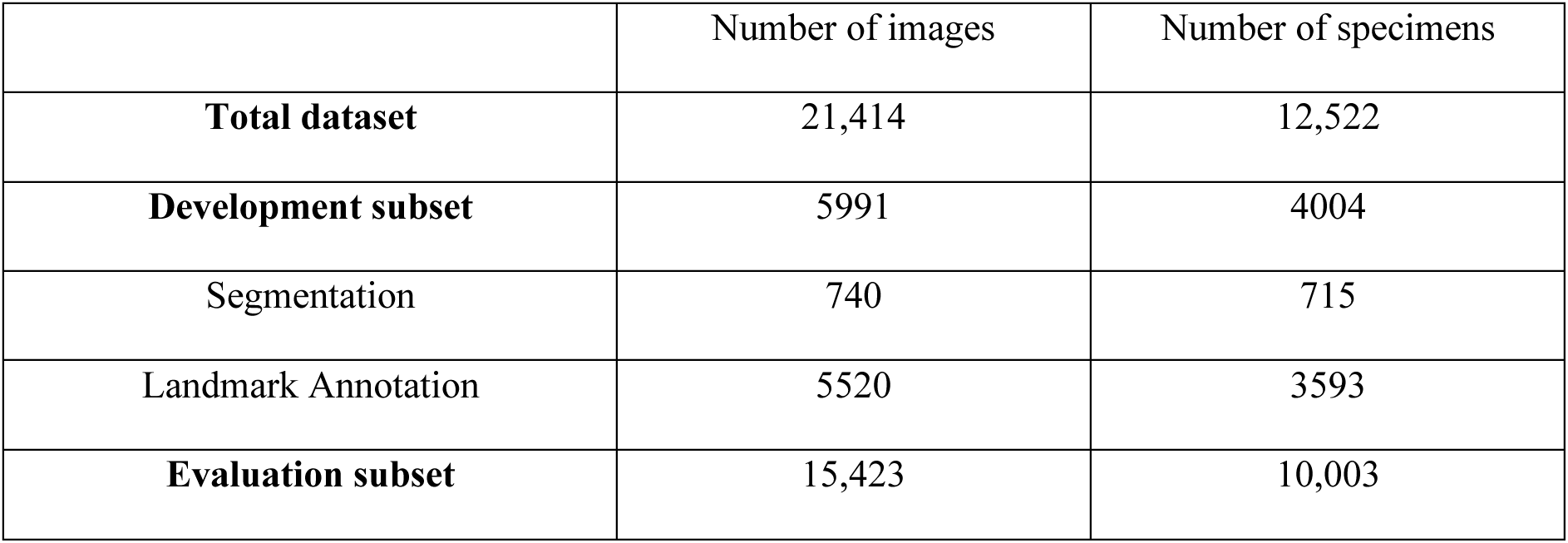
Description of data used to develop and evaluate the landmark annotation system for mosquito wings.

|  | Number of images | Number of specimens |
| --- | --- | --- |
| **Total dataset** | 21,414 | 12,522 |
| **Development subset** | 5991 | 4004 |
| Segmentation | 740 | 715 |
| Landmark Annotation | 5520 | 3593 |
| **Evaluation subset** | 15,423 | 10,003 |

The dataset was split into a “development subset” and an “evaluation subset” (supporting information: metadata_datasplit.csv). The development subset was used to train and validate the UNET++ and the Hourglass models. We provide 5991 annotated images: 740 with ground truth segmentation masks and 5520 with landmark annotations from recent in-house studies, with some images having both annotation types. The evaluation subset contained 12,522 images of undamaged wings from female specimens. These were used to evaluate the system performance after training. There was no overlap between the two subsets.

#### 2.2.2 Model development

Segmentation ground truths were created by generating segmentation masks with napari’s convpaint (Hinderling et al. 2024), followed by manual refinement in Label Studio (https://labelstud.io).

We trained a UNet++ pretrained on ImageNet (Deng et al. 2009; Zhou et al. 2018) using the AdamW optimizer (Loshchilov and Hutter 2019). The loss combined focal loss with skeleton recall loss, a pixel-based loss designed for tubular structures (Kirchhoff et al. 2024). Images were augmented using Albumentations (Buslaev et al. 2018). A five-fold cross-validation scheme was used for both tasks as described by Nolte, Baumbach, et al. (2025). Evaluation metrics were Intersection over Union (IoU) and skeleton recall.

We designed a CoordConv-augmented Hourglass network conditioned on UNet++ segmentation inspired by Wu et al. (2018). The model takes an image and its corresponding segmentation mask as input. The model outputs N landmark heatmaps, where N is the number of desired landmarks. The number of suitable landmark targets depends on the number of homologous junctions on the wing veins. While vein patterns differ between different Dipteran families, within the family they usually share homologous vein junctions (Oosterbroek 2015). For mosquito wings, we adopted a commonly used configuration with 18 landmarks (Champakaew et al. 2021; Sauer et al. 2020; Wilke et al. 2016), but excluded one landmark on the junction of the subcosta and costa as this landmark is difficult to determine and often associated with high observer-dependent variation (Lorenz and Suesdek 2013; Sauer et al. 2026). Thus, our model was trained to annotate 17 landmarks (Figure 2.2).

**Figure 2:**
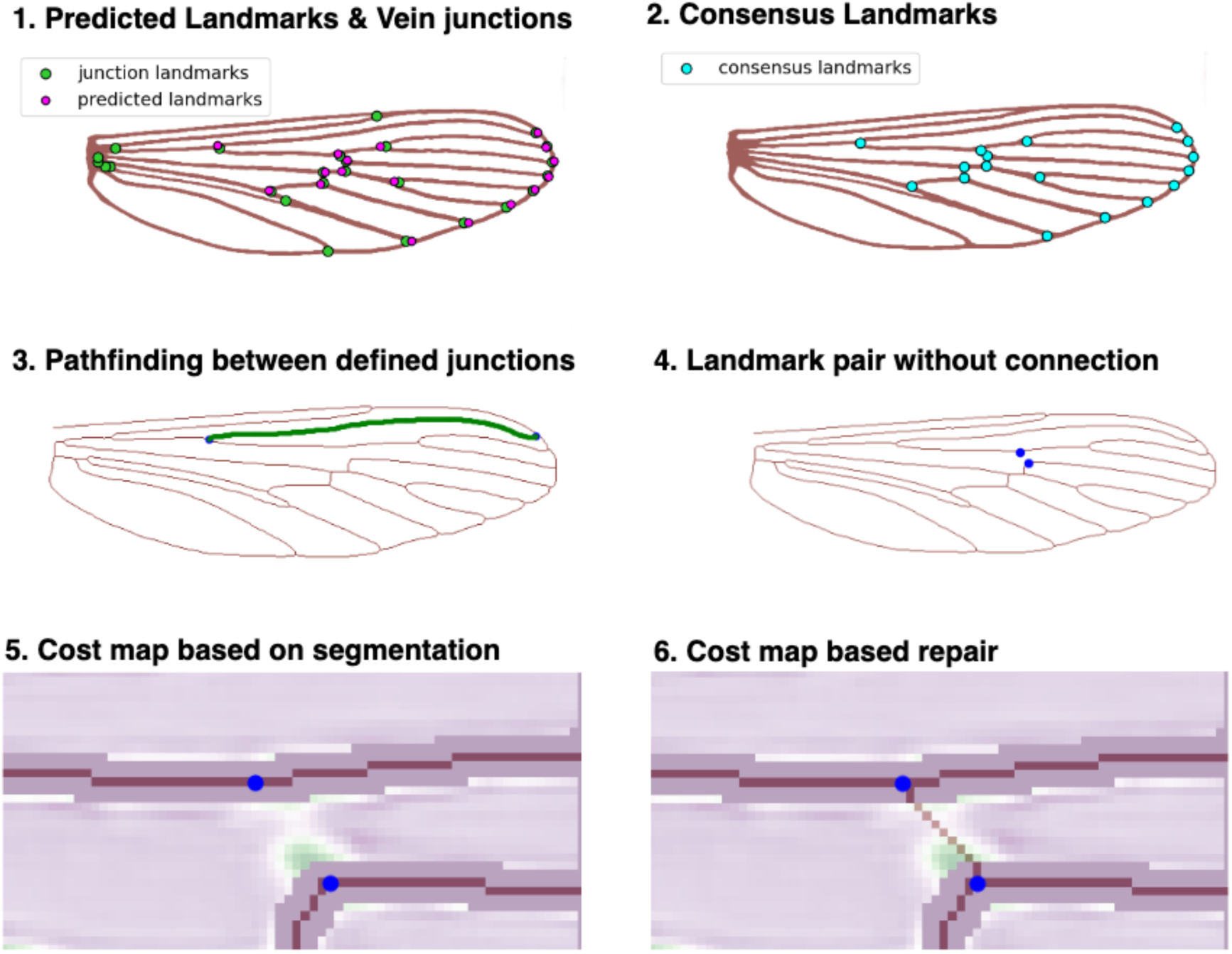
Workflow of the wing representation and repair process. (1) First predicted landmarks and junction landmarks are determined. (2) Consensus landmarks are assigned by finding the closest junction to each predicted landmark. (3) For all defined paths between two points a breath wide search through the wing segmentation is conducted. (4) If a connection could not be created the path is marked as faulty and a repair process is initialized. (5) For wing repair the linear segmentation is used as a cost map (purple: high cost, green: low cost) for a path finding algorithm (6).

Training labels were Gaussian-smoothed landmark heatmaps generated on raw images. These were transformed in lockstep during image preprocessing and geometric augmentations to preserve spatial consistency. Augmentations matched those used for the segmentation model. Additionally, segmentation masks were modified with pixel dropout, coarse dropout, morphological dilation/erosion, and median blurring to simulate plausible UNet++ errors such as small gaps.

The Hourglass model was optimized with AdamW. The loss function combined focal loss and AdaptiveWingLoss (Wang et al. 2020), which were weighted equally. A cost map further weighted the loss by assigning high loss values at the known positions of the other landmarks within each landmark’s prediction layer, discouraging the model from predicting a landmark at a position already occupied by another and thereby reducing the risk of landmark swapping. Landmark coordinates were extracted as argmax positions from the heatmaps. Landmark annotation accuracy was evaluated using mean pixel distance to the human annotated landmarks on the testing folds of each model. For comparison, we benchmarked our system against ML-morph (Porto and Voje 2020). ML-morph uses a combination of object detection and cascade shape regression to predict landmarks. We used the simple-ml-morph implementation (https://github.com/agporto/simple-ml-morph), as the processed images did not benefit from object detection. We applied the same cross-validation splits and evaluated on the same folds used for the Hourglass model.

To contextualise model performance within human annotation variability, we compared the Hourglass model against six human observers, using a subset of 96 images shared with Sauer et al. (2026). Images were standardised to 640×320 pixels using our processing pipeline, with landmark coordinates transformed accordingly. We computed the mean pixel distance to the human annotated landmark for each annotator, the Hourglass model, and ML-morph.

#### 2.2.3 Landmark and Semilandmark Annotation

The final landmark annotation uses both the predicted landmarks and the segmentation. The system aligns the landmarks with the segmentation, as vein junctions in the segmentation did not always match the predicted landmark positions (Figure 2). Each of the 17 landmarks is assigned to its closest vein junction using the Hungarian method, optimising for total distance (Kuhn 1955). After alignment, the system uses breadth-first search to verify the connectivity of the wing vein pattern. A reference dictionary defines allowed connections (where a direct link between two landmarks via a vein is expected) and disallowed connections (where a direct link is not expected and indicates a segmentation error).

When an allowed connection is missing, the system initiated a repair process (Figure 2.5–2.6). This process uses a cost-based function, with the linear output of the segmentation model serving as the cost map. After processing, the wing is re-evaluated for valid or invalid connections and classified as faulty if it is not aligned with the reference dictionary. The complete wing representation (landmarks and their vein connections) is used to place a defined number of semilandmarks at equal fractions of the vein length.

The optimal number of semilandmarks was determined through an experiment using all images from taxa with more than 75 samples the development dataset (*Ae. aegypti, Ae. albopictus, Ae. koreicus, Ae. japonicus, Ae. annulipes-group, Ae. communis-punctor pair, Cx. pipiens* s.l./*torrentium*). We trained LDA models using 5-five-fold cross validation with different number of semilandmarks (N semilandmarks: 0, 11, 21, 30, 41, 52, 63, 74) and varying amount of training samples per taxa labels (4, 6, 8, 10, 12, 14, 16, 18, 20, 25, 30, 35, 40, 45, 50).

#### 2.2.4 Live species identification

To enable species identification via landmark annotation, we randomly selected the landmarks from 100 images per species from all predicted mosquito landmarks as a reference dataset. When a user uploads images, landmarks and semilandmarks are annotated. The predicted landmarks are then merged with the reference dataset, and Generalised Procrustes analysis (GPA) with semilandmark sliding is conducted to align all landmarks. After alignment, the uploaded landmarks are separated from the reference dataset (supplementary information SI1). A LDA model is trained on the aligned reference data and used to predict taxon labels for the uploaded images. Additionally, an CNN model provides a second predictor for classification. The CNN model is developed after Nolte, Baumbach, et al (2025) and retrained using a larger image input size (480×240) and the expanded dataset from Nolte, Agboli et al, (2025). Finally, the results of the two models are merged by a soft-voting scheme.

### 2.3 Experimental Applications

#### 2.3.1 Performance for species identification

To evaluate the automated landmark annotation for species identification, we used the 15,423 images of undamaged wings from female mosquitoes in the evaluation dataset. Taxonomic labels followed Nolte, Baumbach, et al. (2025). Taxa with fewer than 50 samples were grouped into the meta-label “other”, resulting in 34 labels.

We compared the landmark annotation system to a CNN classifier (EfficientNetB0) trained as described in Nolte, Baumbach, et al. (2025). As performance was comparable across all testing folds of the development set, the models evaluated on fold 1 were selected for both tasks (segmentation and landmarking). GPA was applied to align the predicted landmark annotation (Adam et al. 2026; Baken et al. 2021). Five LDA models with covariance matrix shrinkage were trained using five-fold cross-validation, once using landmarks alone and once using landmarks combined with semilandmarks. Sliding semilandmarks were implemented using geomorph (Adam et al. 2026; Baken et al. 2021). Performance was reported as balanced accuracy, and macro F1-score. Five CNN models were trained using the same cross-validation split and performance is reported in the same manner.

#### 2.3.2 Out-of-Distribution Testing

To evaluate the generalisability of ITHILDIN beyond our own data distribution, we reproduced the experiment described by Jeon et al. (2024). In that study, five mosquito species were collected in the Republic of Korea (*Ae. albopictus, An. sinensis, Cx. pipiens* s.s.*, Cx. tritaeniorhynchus, Cx. pallens*) and analysed using wing geometric morphometrics. While the original study distinguished between *Cx. pipiens pallens*, *Cx. pipiens* biotype *pipiens*, and *Cx. pipiens* biotype *molestus*, we excluded *Cx. pipiens* biotype *pipiens* from the reanalysis because these specimens were not collected in the Republic of Korea but in Germany and were part of the training dataset

We replicated the wing geometric morphometric analysis by utilising the application, uploading images to the system and evaluating the returned prediction. We compared the performance of the CNN-based prediction, the landmark-based prediction and the ensemble-based prediction. Not all taxa labels in Jeon et al. (2024) are identical to ours. We defined a successful prediction when *Ae. albopictus* was predicted as *Ae. albopictus* taxa label, *Cx. pipiens* biotype *molestus* and *Cx. pallens* were expected as the *Cx. pipiens* s.l./*torrentium* label and *Cx. tritaernorhynchus* was expected as *Cx. vishnui*-group. We had no trained taxa label for *An. sinensis* therefore we evaluated performance on it separately.

#### 2.3.3 Cross-Family Transferability

To demonstrate the transferability of ITHILDIN to other families, we adapted it to two additional Diptera families: Drosophilidae and Glossinidae each represented by the genus *Drosophila* and *Glossina* respectively.

For Glossinidae, we used the dataset from Geldenhuys et al. (2023) (n = 2421 images) with additional images (n = 320) from Nolte, Agboli, et al. (2025). For segmentation retraining, we manually labelled 116 wing images (57 from Geldenhuys et al.; 59 from Nolte, Agboli, et al.) following the protocol described in Section 2.2.2. The segmentation model was trained using the same procedure as for mosquitoes, initialised with pretrained weights. For landmark annotation, we processed all images from Geldenhuys et al. (2023) and generated corresponding landmark heatmaps. Training followed the protocol for mosquitoes with an 80:20 train–test split, again using pretrained weights. We evaluated mean pixel distance on the test set. To assess classification performance, we used the 257 images from Nolte, Agboli, et al. (2025), excluding all samples which were used to train the segmentation model. The test set contained three taxa: *G. fuscipes* s.l. (n = 101), *G. palpalis gambiensis* (n = 86), and *G. palpalis palpalis* (n = 70). We trained and evaluated LDA models using five-fold cross-validation.

For Drosophilidae, we used the dataset (n = 1134) from Sonnenschein et al. (2015) evaluating classification strategies of *Drosophila melanogaster* wings, accessed via Nguyen et al. (2022). For segmentation training, we sampled 29 images and created ground truth segmentation maps following the protocol described above for mosquitoes. For landmark training, we used the provided landmark annotation. Models were trained and evaluated using five-fold cross-validation. Mean pixel distance was assessed on the combined test folds across all folds. To assess classification performance, we trained LDA models on the predicted landmarks using five-fold cross-validation to discriminate between four mutant genotypes present in the dataset.

## 3 Results

### 3.1 Pipeline Development

#### 3.1.1 Model development

For the development of the segmentation model, five models were trained using five-fold cross-validation. Performance is reported on the four folds used exclusively for testing. Segmentation models achieved a mean skeleton recall of 85.2% (95% CI: 84.6–85.7), a mean IoU of 70.7% (95% CI: 70.2– 71.1), and a soft Dice loss of 0.1887 (95% CI: 0.1866–0.1908).

For landmark annotation the mean pixel distance between ground truth and prediction was 2.8 pixels (95% CI: 2.7–2.9) on an image resolution of 640×320 (Figure 3) (supporting information: landmark_hourglass_prediction.csv). Landmark 4 was predicted best with a mean pixel distance of 2.4 (95% CI: 2.3–2.5) whereas landmark 8 had the highest mean pixel distance at 4.2 (95% CI: 4.0–4.5).

**Figure 3:**
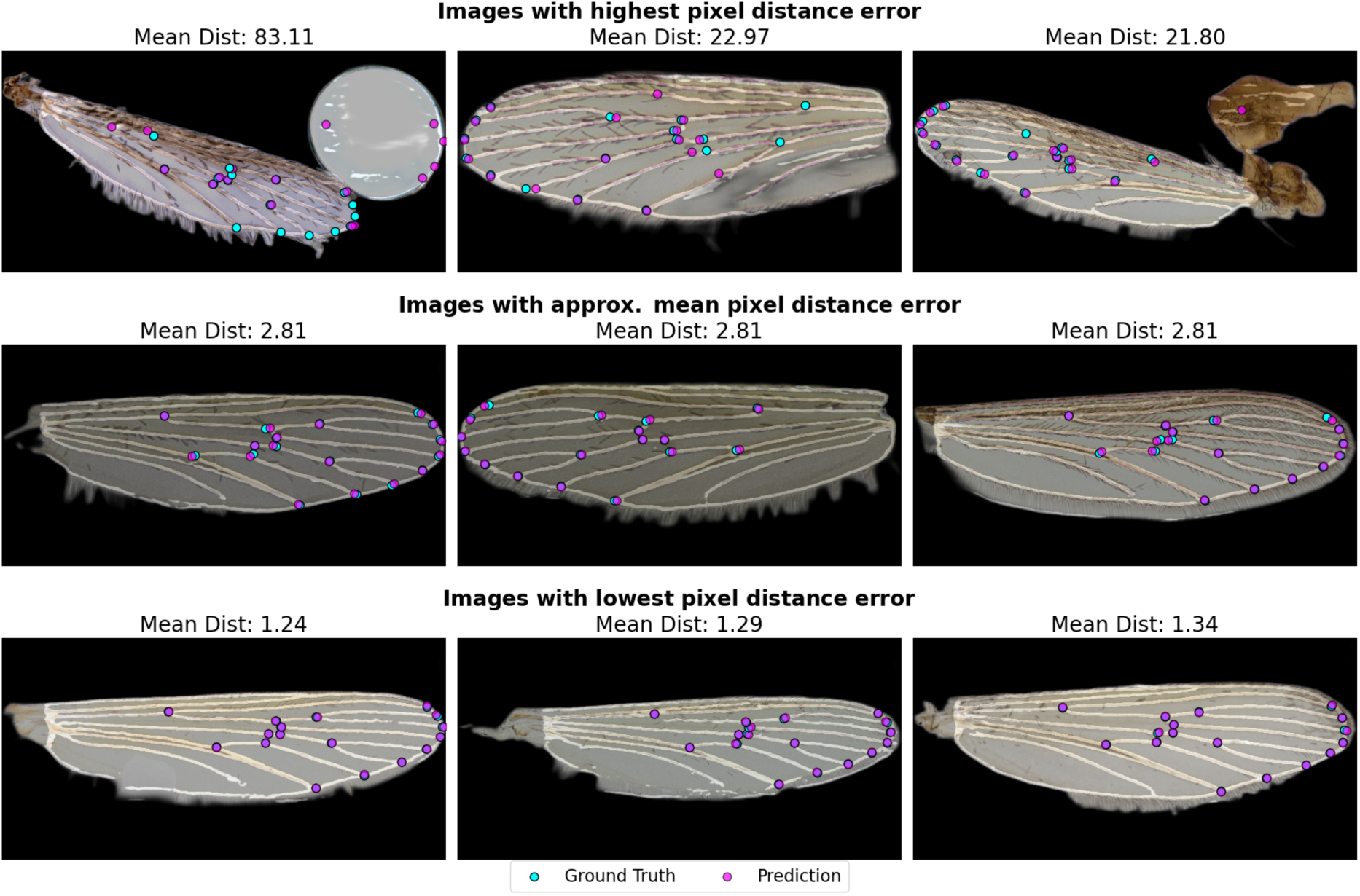
Examples wing images with predicted landmarks (purple) compared to manually annotated landmarks (cyan). Top row shows the 3 images with highest mean pixel distance to the ground truth. Middle row shows images with mean performance values and bottom row present images with lowest mean pixel distance.

We compared performance to ML-morph under the same training and testing configuration. Mean pixel distance was 10.7 (95% CI: 9.9 - 11.4) on the same image resolution (supporting information SI2, landmark_mlmorph_prediction.csv). Landmark 13 was predicted with smallest mean pixel distance at 9.0 (95% CI: 8.6 - 9.4). Landmark 16 had the highest mean pixel distance at 13.5 (95% CI: 12.8 – 14.2).

To contextualise the error of the landmark model to human observer variability we compared its performance on a subset of images which were also annotated by six different human observers in Sauer et al (2026). The Hourglass model achieved a mean pixel distance of 4.5 pixels (95% CI: 4.3–4.6), to ground truth, placing it within the range of human observer variability across the six annotators 4.7 (95% CI 4.4 - 5.0). ML-morph yielded a mean pixel distance of 12.7 (95% CI: 11.1 – 14.2).

#### 3.1.2 Landmark and Semilandmark Annotation

The system uses predicted landmarks and segmentation to construct a wing representation for semilandmark placement. Of the 15,423 samples in the evaluation dataset, 43.7% had a complete wing representation without the need to repair the segmentation map. The remaining samples underwent the repair process: 43.9% were successfully repaired, while 12.4% could not be repaired or contained disallowed connections.

Failure rates (unsuccessful repairs) varied by genus and imaging device (supplementary information SI3). The smartphone with macro lens had the highest failure rate (28.8%), whereas images captured with a microscope had a considerably lower failure rate (Olympus SZ61 & Olympus DP23: 11.0% and Leica M205C 11.9%). Among the three most represented genera within the dataset, *Anopheles* had the highest failure rate (25.9%), while *Aedes* and *Culex* failed at 15.4% and 2.8%, respectively.

To determine the optimal number of samples and semilandmarks for classification, we ran an optimisation experiment using the most represented taxa from the development dataset. Classification accuracy increased consistently with the number of samples per class across all semilandmark configurations, rising from 80% at 4 samples to above 90% with 25 samples, with diminishing returns thereafter (Figure 4). Generally, higher amounts of semilandmark consistently yielded higher balanced accuracy up to a certain point. with the best-performing configuration (52 semilandmarks) reaching 95.0% at 50 samples per class.

**Figure 4:**
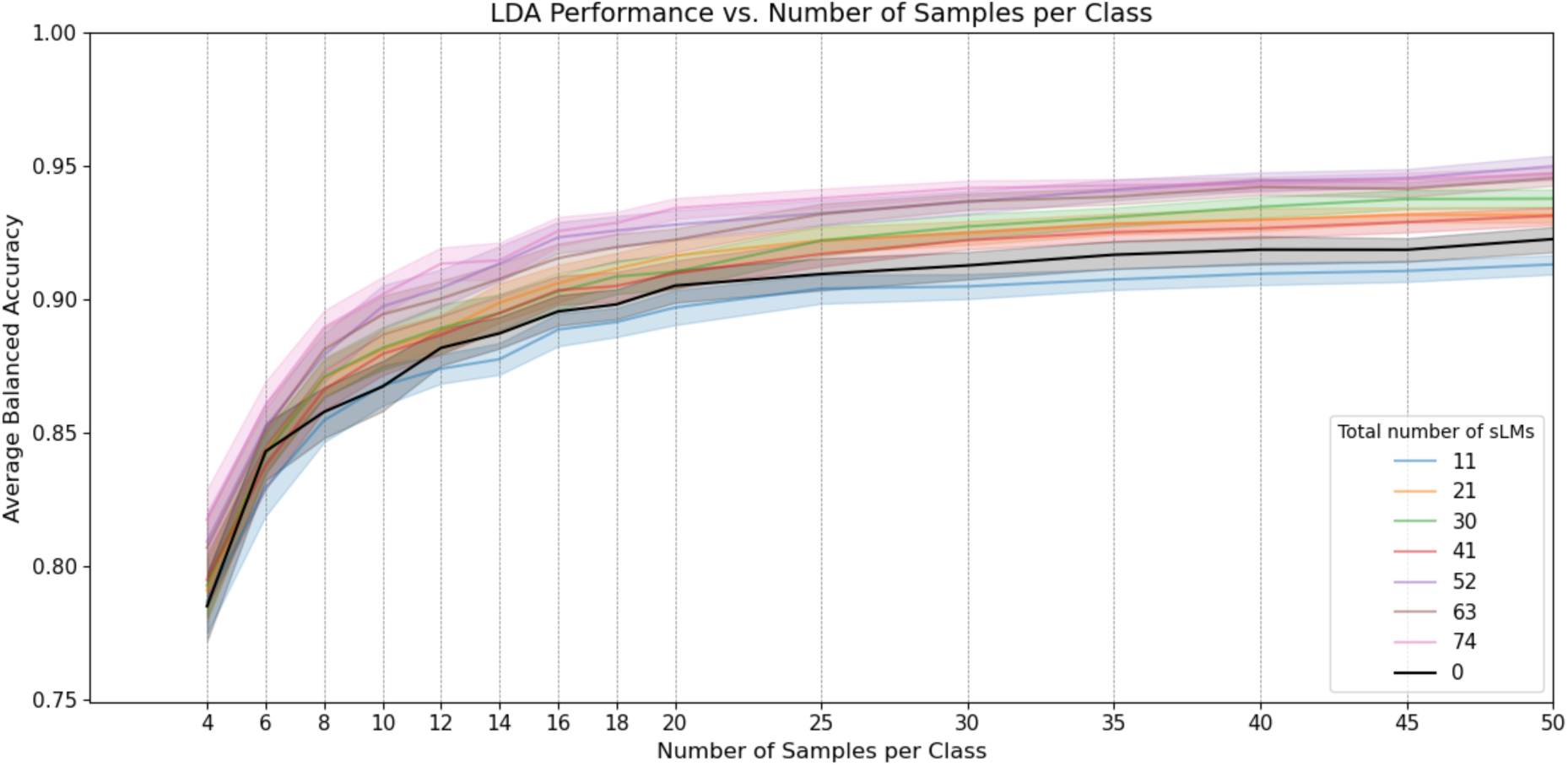
Optimization experiment where different combinations of number of semilandmarks and number of training samples were tested for performance utilising the development dataset.

### 3.2 Experimental Applications

#### 3.2.1 Performance for species identification

We benchmarked the landmark annotation system against ML-morph and a CNN by comparing the taxonomic classification accuracy. We evaluated all methods under two imaging conditions: microscope images only, and microscope and phone images combined. Furthermore, we report results for two sample subsets: all samples, including those where semilandmark annotation failed, and non-failed samples only (supporting information: landmark_ITHILDIN_speciesclass.csv).

For the non-failed samples, the CNN classifier and semilandmark approach achieved the highest balanced accuracies at 94.0% and 90.8%, respectively (F1-scores: 92.8% and 88.8%) when evaluated on the full dataset (Table 2). The 17 landmark approach produced lower performance on the full dataset with 75.1% (CI95%: 73.3 - 77.0) balanced accuracy but improved in performance when restricted to the non-failed subset (Balanced Accuracy: 86.0% (CI95%: 84.6 - 87.3)) The landmarks based on the ML-morph annotation resulted in the worst classification performance where performance measured either as balanced accuracy or macro F1-Score was around 22%. All models showed by far the lowest performance on the “other” class, with accuracy ranging around 57% for the CNN model and down to 12% for the landmark LDA models.

**Table 2:** Performance of different landmark annotation methods and CNN classifier in distinguishing between 34 Taxa of the evaluation dataset.

| Mode | Device | Metric | Balanced Accuracy<br>(95% CI) | F1 Score<br>(95% CI) |
| --- | --- | --- | --- | --- |
| CNN Classifier | Phone &<br>Microscope | all | 94.2 (92.6 - 95.8) | 93.0 (91.5 - 94.5) |
|  |  | non-failed | 94.0 (91.7 - 96.2) | 92.8 (90.8 - 94.9) |
|  | Microscope | all | 94.1 (92.6 - 95.6) | 93.0 (91.7 - 94.3) |
|  |  | **non-failed** | 93.9 (91.6 - 96.2) | 92.8 (90.7 - 94.8) |
| **ITHILD Landmark** | **Phone &**<br>**Microscope** | **all** | 75.1 (73.3 - 77.0) | 74.0 (72.8 - 75.2) |
|  |  | **non-failed** | 86.0 (84.7 - 87.3) | 84.1 (82.7 - 85.4) |
|  | **Microscope** | **all** | 78.2 (77.2 - 79.2) | 76.1 (75.3 - 76.9) |
|  |  | **non-failed** | 86.1 (84.6 - 87.6) | 84.2 (82.6 - 85.8) |
| **ML-morph**<br>**Landmark** | **Phone &**<br>**Microscope** | **all** | 22.1 (21.7 - 22.5) | 22.7 (22.2 - 23.1) |
|  |  | **non-failed** | 22.9 (22.4 - 23.5) | 23.2 (22.5 - 24.0) |
|  | **Microscope** | **all** | 21.5 (21.2 - 21.9) | 21.9 (21.6 - 22.2) |
|  |  | **non-failed** | 22.3 (21.5 - 23.1) | 22.5 (21.4 - 23.6) |
| **ITHILDIN**<br>**Semilandmark** | **Phone &**<br>**Microscope** | **non-failed** | 90.8 (89.6 - 92.1) | 88.8 (87.5 - 90.2) |
|  | **Microscope** | **non-failed** | 90.8 (89.1 - 92.4) | 89.0 (87.3 - 90.7) |

#### 3.2.2 Out-of-Distribution Testing

To evaluate the system’s performance on out-of-distribution data, we replicated the study from Jeon et al. (2024) by utilising the landmark annotation and species prediction of the ITHILDIN application (supporting information: landmark_ITHILDIN_jeon.csv). Prediction results solely based on the CNN resulted in moderate success. Balanced accuracy for known classes (*Cx. pipiens* biotype *molestus, Cx. pallens, Cx. tritaernorhynchus, Ae. albopictus*) was 66.1%. While *Cx. pipiens* biotype *molestus* and *Cx. pallens* were identified with over 80% accuracy, the model showed limited ability to classify, *Cx. tritaernorhynchus* (29%) and distinguish between *Ae. albopictus* and *Ae. aegypti,* only achieving an accuracy of 50% while predicting *Ae. aegypti* in 42% of cases (Figure 5). Using landmarks and semilandmarks, the system achieved a balanced accuracy of 77.3%. The failure rate to annotate semilandmarks was 9.4%. Here we observe the lowest classification performance for *Cx. pallens* (66%) and *Cx. tritaernorhynchus* (71%). Best performance was observed when using the soft-voting ensemble where we observe a balanced accuracy of 87.6% on the known taxa.

**Figure 5:**
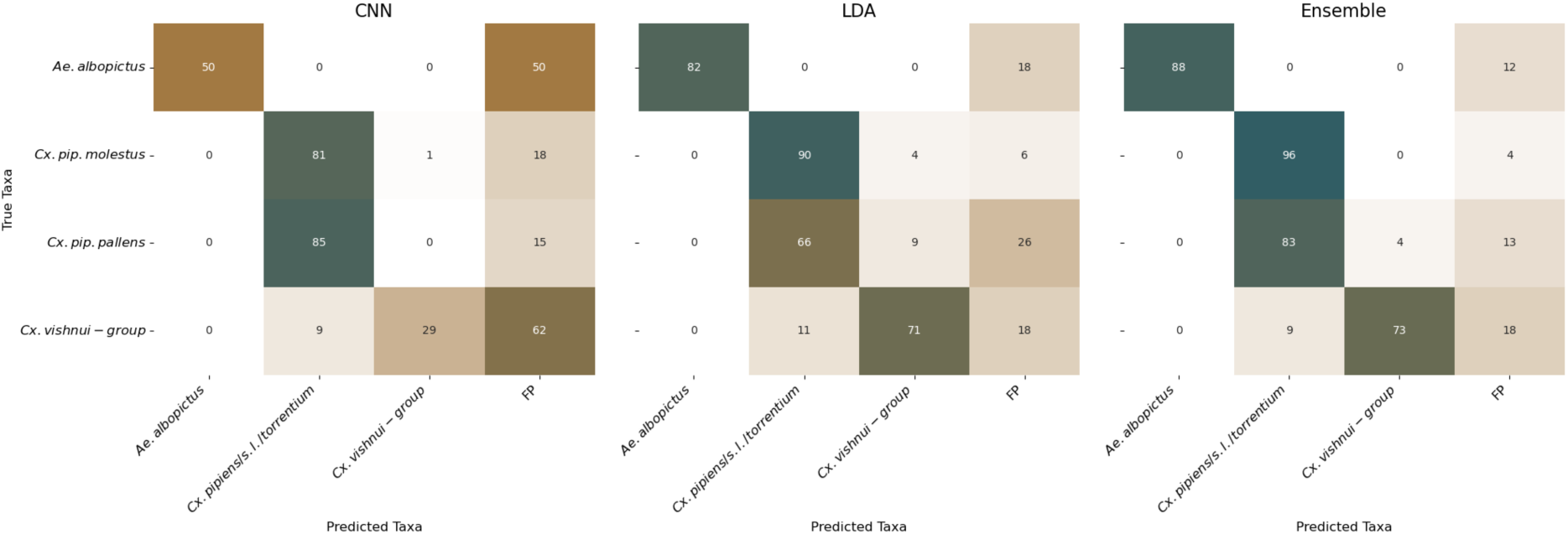
Accuracy of different classification methods (CNN, LDA, soft-voting Ensemble) on out-of-distribution data from Jeon et al (2024).

When replicating the study from Jeon et al. (2024) by training a LDA model on the predicted landmarks in a five-fold cross-validation split we report a balanced accuracy of 94.8% (CI95%: 92.7 - 96.9) utilising landmarks and when including semilandmarks we report 97.7% (CI95%: 95.8 - 99.6) balanced accuracy. This is comparable to the performance reported by Jeon et al. (2024) on their manual landmark annotations.

#### 3.2.3 Cross-Family Transferability

For Glossinidae, the retrained system achieved a mean pixel distance of 5.0 pixels (CI95%: 4.8–5.3) on an image resolution of 640×320 (supplementary information SI4). For classification of the three *Glossina* taxa using LDA, the landmark-only approach achieved a balanced accuracy of 69.1% (95% CI: 62.9–75.4) and F1-score of 69.3% (95% CI: 63.0–75.6). Performance improved substantially with semilandmarks, reaching a balanced accuracy of 83.3% (CI95%: 78.4-89.7) and F1-score of 83.2% (CI95% 77.5 - 88.9).

The system achieved a mean pixel distance of 3.7 pixels (95% CI: 3.5–3.8) across all Drosophilidae landmarks on images in 640×320 resolution (supplementary information SI5). For classification of the four *Drosophila* loss-of-function mutations, the landmark-only approach achieved a balanced accuracy of 60.3% (95% CI: 56.8–63.8) and F1-score of 60.2% (95% CI: 56.8–63.5). Performance improved with semilandmarks, reaching a balanced accuracy of 71.1% (95% CI: 65.6–76.6) and F1-score of 71.0% (95% CI: 65.8–76.3).

## 4 Discussion

The objective of this study was to make wing geometric morphometrics of Diptera more reproducible and accessible using computer vision. The result is a publicly available system (https://ithildin.bnitm.de) that automates the entire landmark and semilandmark annotation process and applies it for species identification.

The developed system annotates landmarks reliably, with marginal pixel error, performing within the variance expected between human observers and outperforming other state-of-the-art automated landmark annotation techniques namely ML-morph. However, classification performance using the 17 landmarks and LDA varied with task complexity. In small-scale experiments with few taxa, performance was strong, with the relatively straightforward five-species distinction in Jeon et al. (2024) achieving 94% balanced accuracy. When applied to the larger and more challenging evaluation dataset of 34 taxa, including multiple morphologically similar species pairs, landmark-only classification yielded 75% balanced accuracy (86% when excluding annotation failures), suggesting that 17 landmarks alone may provide insufficient shape resolution for reliable discrimination in a more taxonomically diverse context. This is consistent with the theoretical expectation that morphological characterisation improves with an increasing number of landmarks (Watanabe 2018), and is further supported by our optimisation experiment, in which increasing semilandmark density generally improved species classification accuracy. On the 34-taxa evaluation dataset, the semilandmark-based approach achieved 5% increase in balanced accuracy (91%), approaching the CNN classifier (94%). The most pronounced improvement was observed for the morphologically similar Glossinidae taxa, where landmark-only classification achieved 69%, and the addition of semilandmarks yielded a 14% absolute improvement to 83%. These results demonstrate the value of semilandmarks for taxonomic classification tasks involving diverse or morphologically similar taxa.

The system’s applicability is constrained by a failure rate of approximately 12%. While the 17 landmarks could theoretically still be used when semilandmark placement fails, the inability to construct a valid representation of the wing typically indicate that either segmentation or landmark prediction produced an unreliable result. Failure rate was associated with image quality. Images captured with a smartphone had the highest failure rate (29%), whereas microscope-captured images failed less (11-12%). We hypothesize that certain wing characteristics also increased failure, such as the dark wing scale spots typical for certain mosquito taxa e.g., *An. maculipennis* s.l. or *Culiseta annulata*. Additional training data may address the issue, as the current segmentation dataset is dominated by samples from *Culex* and *Aedes* mosquitoes. However, image quality remains paramount, and we do not recommend using the system with smartphone-captured images. Yet the ability to recognize the failure serves as an implicit quality control mechanism as seen by the increase of performance for species identification via 17 landmarks when failed samples are removed.

Generalisation to out-of-distribution data is a known weakness of deep learning classifiers (D’Amour et al. 2022) and the CNN’s limited performance on the Jeon et al. (2024) dataset is consistent with this, despite applying mitigation techniques recommended by Nolte, Baumbach et al. (2025). The landmark-based classifier performed considerably better, which is expected given that geometric shape representations are less vulnerable to domain shift than pixel-level CNN features. However, the landmark classifier still performed notably worse in the live experiment than in the closed experiment, suggesting another limiting factor. In the live application, specimens are classified by comparison to a reference dataset. Our reference dataset contains landmarks for all taxa within the *Cx. pipiens* s.l./*torrentium* complex except *Cx. pallens*, meaning that *Cx. pallens* specimens could only be assigned by similarity to the other taxa rather than to a dedicated reference. Yet, we observe that *Cx. pallens* is morphologically distinct and discriminable from its sibling taxa by its successful classification in the closed experiment. This suggests that the failure reflects a gap in the reference data rather than a limitation of the landmark prediction itself. Like molecular identification methods, the landmark classification system is therefore bounded by the completeness of its reference data (Fujisawa and Imai 2026). Expanding this reference is, however, considerably more accessible than building molecular databases, requiring only image acquisition.

Beyond taxonomic classification, the system offers analytical value that distinguishes it from purely CNN-based approaches. Because it is grounded in geometric morphometrics, it provides an interpretable and biologically meaningful representation of wing shape, enabling analyses beyond identification. For instance, the same landmark and semilandmark coordinates can be used to investigate intraspecific variation, such as morphological differences across geographic populations or in response to environmental factors (Lorenz et al. 2017). The system also demonstrated that fine-grained morphological distinctions are achievable as it successfully distinguished *Cx. pallens* from *Cx. pipiens* biotype *molestus* (Jeon et al. 2024). Together, these properties position the system not just as a classification tool, but as a flexible and interpretable tool for geometric morphometric research.

Although developed on mosquitoes, the system transferred easily to other Diptera families. The modular pipeline architecture enabled successful adaptation to Drosophilidae and Glossinidae. Landmark annotation accuracy remained around or below 5 mean pixels error for both families, comparable to mosquitoes. In addition, we observe similar classification performance for Drosophilidae as reported by Sonnenschein et al. (2015).

A key advantage of automated annotation is its value as a consistent, single observer for landmark placement. As suggested by Dujardin et al. (2010) standardising annotations to a single observer provides a viable strategy to combine morphometric datasets across studies and research groups. By eliminating inter-observer variability, the system enables large-scale analyses that would otherwise be confounded by observer-specific biases. This is further supported by the mostly untapped potential of existing morphometric data in the literature: landmark coordinates and wing images from published studies represent a substantial resource that, combined with more widespread open-access data sharing, could be leveraged for large-scale analyses using this system as a standardising tool.

This study demonstrates that integrating geometric morphometrics with computer vision provides a promising approach for Diptera wing analysis. The accuracy of automatic landmark annotation was comparable to that of human observers. Species classification based on these automatically obtained landmarks and semilandmarks performed similar to CNN classifiers, while providing advantages over purely classifier-based approaches: interpretable shape representations, reliable failure detection, and the flexibility to support broader morphometric analyses such as population-level or environmental variation studies.

## Supporting information

contains supporting information

## Acknowledgements

The authors would like to thank the Federal Ministry of Research, Technology and Space of Germany (BMFTR) under the project NEED and CIMT (Grant Number 01Kl2022). FGS received funding of the Klaus Tschira Boost Fund, a joint initiative of GSO – Guidance, Skills & Opportunities e.V. and Klaus Tschira Stiftung. This project has received resources from the Infravec consortium as part of the ISIDORe project (funding from the European Union’s Horizon Europe Research & Innovation program, grant agreement N° 101046133).

Generative artificial intelligence tools were used in a limited and supportive capacity during the preparation of this manuscript. Specifically, such tools were employed to assist with improving the clarity and language of the text and to support the development and documentation of code used. These tools were not used to generate scientific content, interpret results, or draw conclusions. All outputs were reviewed, verified, and edited by the authors, who take full responsibility for the accuracy, integrity, and originality of the work.

The authors declare no conflict of interest.

## Data availability statement and Data sources

Images used in this study are accessible under CC BY 4.0 license at https://doi.org/10.6019/S-BIAD1478. Downloadable and installable docker application can be accessed on the applications’ git page: https://anonymous.4open.science/r/ITHILDIN-4313/. A demonstrator of the application is hosted on https://ithildin.bnitm.de/

## Author contributions

KN, FGS, RL, PK and JB conceived the ideas and designed methodology; KN, FGS and RL collected the data; KN, FGS and RL analysed the data; KN, FGS and RL led the writing of the manuscript. All authors contributed critically to the drafts and gave final approval for publication.

