## Supplementary material for "Automated landmark and semilandmark annotation for wing geometric morphometrics in Diptera using deep learning": contains supporting information

#### SI1: Workflow of ITHILDINs Live Prediction

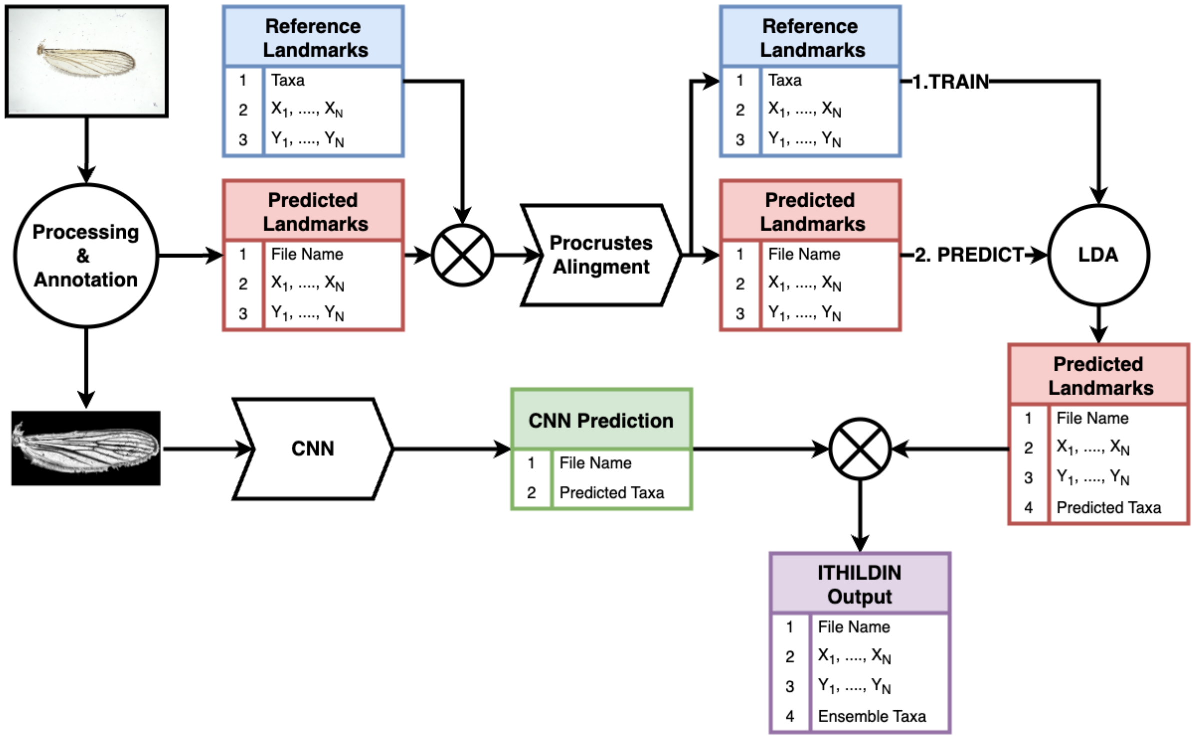


*Figure 1:Schematic workflow of ITHILDINs live species prediction system. After predicting landmarks for the uploaded image, they are merged with a reference dataset for a Procrustes alignment. Hereafter they are separated, and the reference landmarks are used to train a LDA model which then predicts the species for the aligned predicted landmarks. A CNN is used to classify the processed image and both predictions are then merged in a soft-voting ensemble to produce the final prediction.*

### SI2: Mean Pixel Distance examples ML-morph

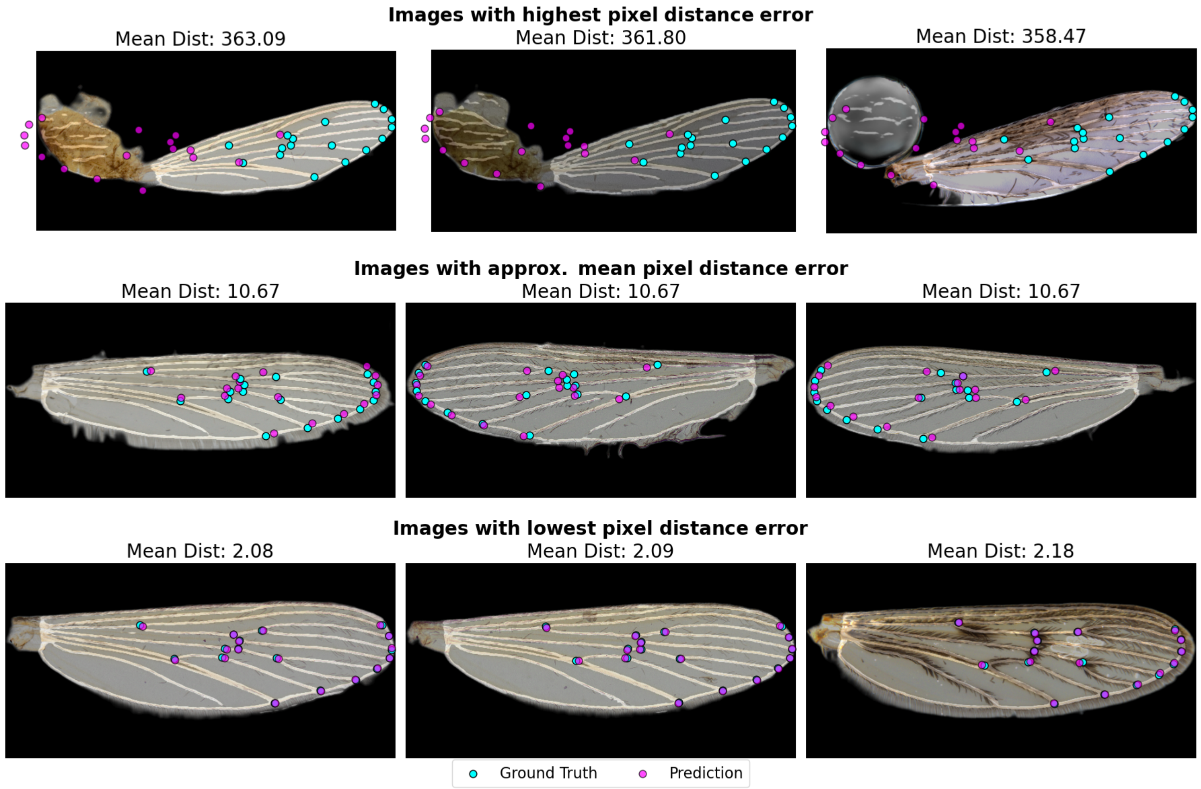


*Figure 2: Examples wing images with predicted landmarks from ML-morph (purple) compared to manually annotated landmarks (cyan). Top row shows the 3 images with highest mean pixel distance to the ground truth. Middle row shows images with around mean performance values and bottom row presents images with lowest mean pixel distance.*

### SI3: Distribution of Failure Rate per genus, taxa label and device

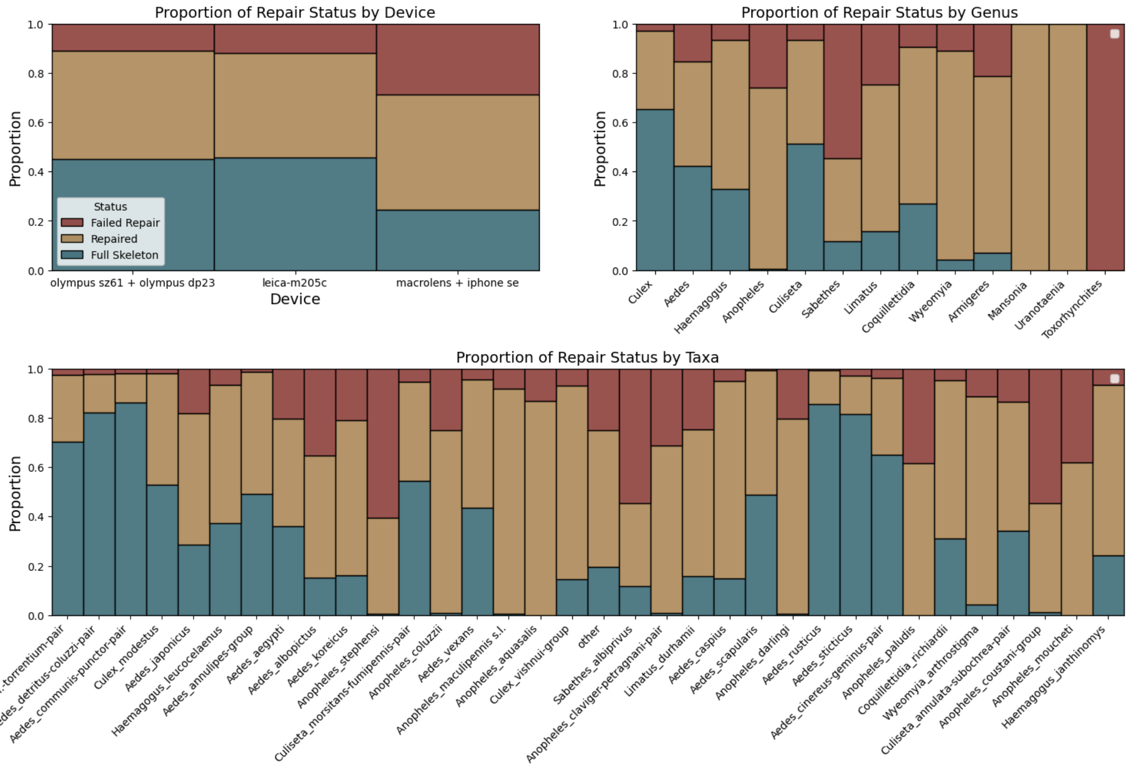


*Figure 3: Distribution of Failure Rate per genus, taxa label and device.*

#### SI4 mean pixel distance examples *Glossina*

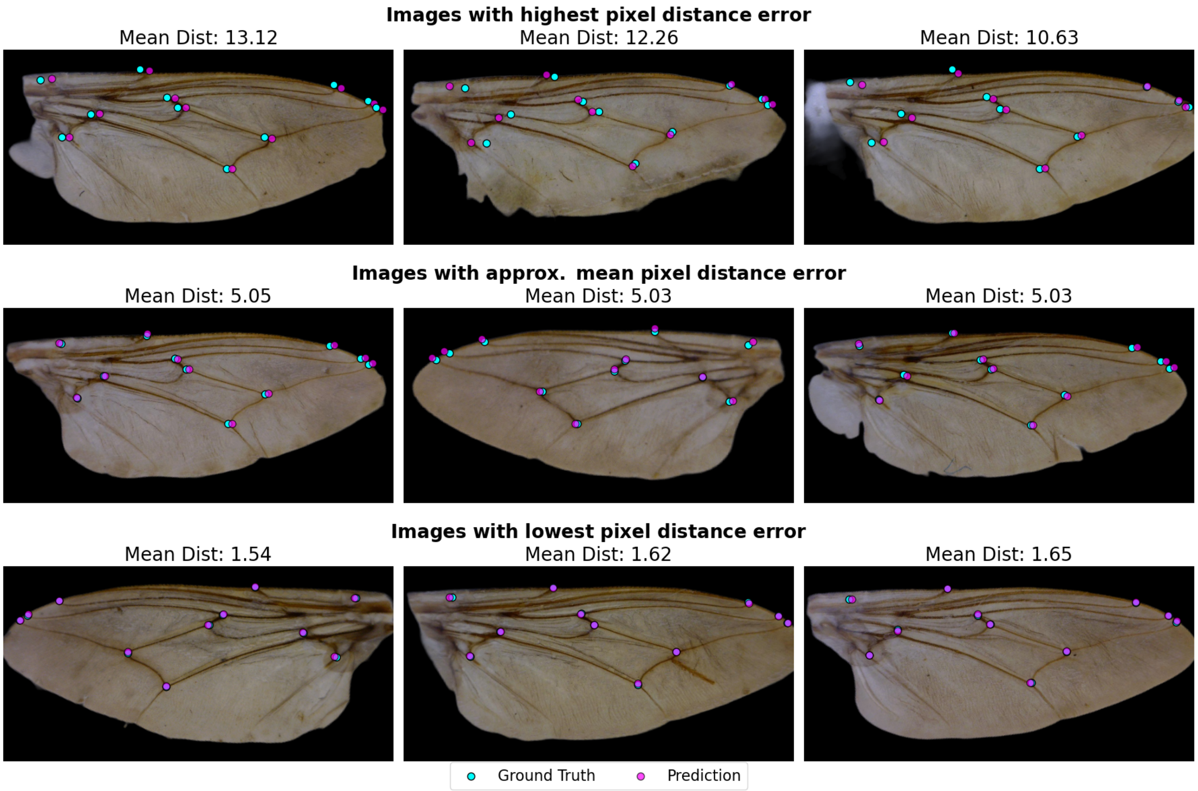


*Figure 4: Examples Glossina wing images with predicted landmarks (purple) compared to manually annotated landmarks (cyan). Top row shows the 3 images with highest mean pixel distance to the ground truth. Middle row shows images with around mean performance values and bottom row presents images with lowest mean pixel distance.*

### SI5 Mean pixel distance examples *Drosophila*

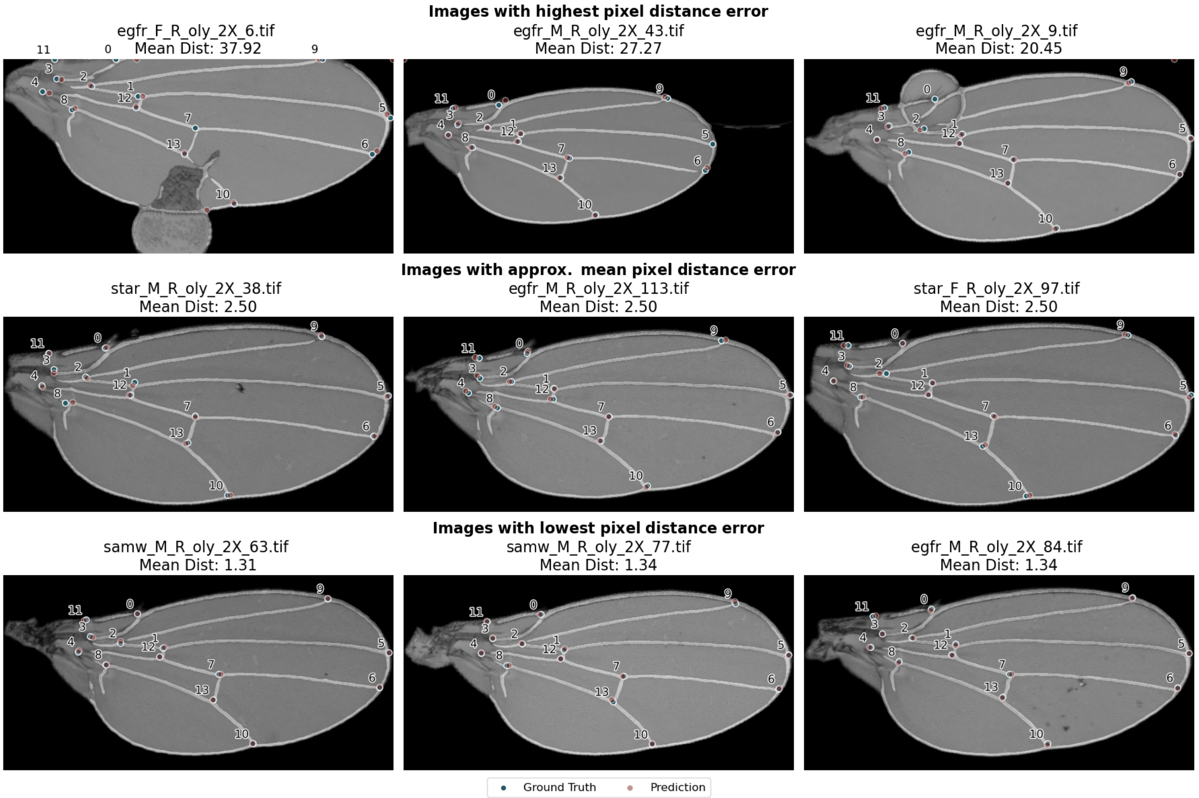


*Figure 5: Examples Drosophila wing images with predicted landmarks (purple) compared to manually annotated landmarks (cyan). Top row shows the 3 images with highest mean pixel distance to the ground truth. Middle row shows images with around mean performance values and bottom row presents images with lowest mean pixel distance.*
